# Biological Boundary Conditions Regulate the Internalization of *Aspergillus Fumigatus* Conidia by Alveolar Cells

**DOI:** 10.1101/2024.11.12.623156

**Authors:** Natalia Schiefermeier-Mach, Julien Polleux, Lea Heinrich, Lukas Lechner, Olexandra Vorona, Susanne Perkhofer

## Abstract

**Introduction:** The lung environment is defined by unique biological boundary conditions, including complex alveolar geometry, extracellular matrix composition and mechanical forces generated during respiration. These factors were shown to regulate alveolar permeability, surfactant secretion, cell contractility and apoptosis, but their role in fungal infections remains unknown. *Aspergillus fumigatus* is a critical fungal pathogen that causes severe pulmonary infections in immunocompromised individuals. Our study addresses a knowledge gap by investigating how boundary conditions affect *A. fumigatus* conidia interactions with alveolar epithelial cells.

**Methods:** We applied micropatterned substrates to confine cells into defined shapes and densities, allowing precise control over geometric conditions and extracellular matrix composition. Using cell line stably expressing the phagolysosomal protein Lamp1-NeonGreen and multiplane fluorescent microscopy, we evaluated *A. fumigatus conidia* binding and internalization efficiency.

**Results:** We observed significantly faster and more efficient *A. fumigatus* conidia internalization in cells confined on micropatterns compared to previously reported studies using cell monolayers. Altering cell geometry, density, and extracellular matrix composition strongly affected conidia binding and localization to Lamp1^+^ phagolysosomes. Cells on X-shaped or multicellular micropatterns showed higher internalization rates, particularly at the periphery, suggesting spatial heterogeneity in pathogen uptake. Additionally, changes in extracellular matrix composition influenced the intracellular trafficking of *A. fumigatus* conidia.

**Discussion:** Our findings emphasize the essential role that local mechanical and biochemical cues play in shaping the interactions between fungal pathogens and alveolar cells. Understanding how lung boundary conditions change in disease states will provide important insights into fungal infection outcomes.

## 1 Introduction

Biological boundary conditions are the physical and biochemical constraints that define the cellular environment and regulate cellular responses. These conditions include cell number, shape, size, extracellular matrix (ECM) composition, substrate stiffness and intercellular forces (Vahey and Fletcher, 2014). Investigating boundary conditions is important for understanding how cells interact with their environment and how these interactions influence their function. They have been shown to regulate essential cellular processes, such as adhesion, migration, proliferation, endocytosis, and bacteria-cell binding (Wickström et al., 2010; Hamidi et al., 2020; Bastounis et al., 2022; Wang et al., 2022; Feng et al., 2023; Isomursu et al., 2024).

Micropatterned substrates have provided important insights into how geometric constraints affect cellular functions (Azioune et al., 2010; Muoth et al., 2016; Wickström and Niessen, 2018; van der Putten et al., 2022). Confined cells to specific shapes alter their biomechanical properties, including cytoskeletal organization and focal adhesion dynamics, consequently influencing intracellular signaling pathways (Schauer and Goud, 2014; He et al., 2015). For example, cells on circular micropatterns exhibit distinct polarization patterns regulated by mechanical forces, like shear stress and tension (He et al., 2015). Schauer and Goud (Schauer and Goud, 2014) showed that geometric constraints alter endocytic activities, leading to asymmetric uptake of transferrin and epidermal growth factor ligands. Biological boundary conditions have been also suggested to determine the fate of internalized molecules and microorganisms (Muoth et al., 2016; Bastounis et al., 2022).

The role of boundary conditions is particularly relevant in host-pathogen interactions, where they can regulate pathogen survival within tissues. Previous research has shown that bacterial pathogens such as *Listeria monocytogenes* and *Shigella flexneri* exploit host-cell mechanics to spread by manipulating mechanical forces at cell-cell junctions and altering intracellular tension (Bastounis et al., 2022). Furthermore, Feng et.al recently showed that the initial adhesion of *Staphylococcus aureus* and *Escherichia coli* to host cells can be modulated by altering ECM rigidity and geometric constraints (Feng et al., 2023).

While studies of bacterial models have been investigated, the impact of boundary conditions on fungal pathogens remains unexplored. Fungal infections cause over 3.8 million deaths annually, with *Aspergillus* species responsible for the highest mortality rates among fungal diseases (Denning, 2024). *Aspergillus fumigatus* is recognized as one of the top four critical fungal pathogens by the WHO (WHO Antimicrobial Resistance Division, 2022). These saprotrophic fungi produce dense clouds of conidia containing up to 10^8^ spores per cubic meter. In healthy individuals, conidia are cleared in the airway by innate immune mechanisms including mucociliary clearance and macrophage phagocytosis (Latgé, 2001). However, in immunocompromised individuals, compromised immune defenses allow *A. fumigatus* to survive in the lungs, causing various forms of pulmonary aspergillosis, ranging from mild hypersensitivity to severe invasive infections (Dagenais and Keller, 2009; Vanderbeke et al., 2018; Latgé and Chamilos, 2019).

Interactions between lung cells and *A. fumigatus* conidia is a complex process that involves both professional phagocytic cells and non-professional phagocytes like alveolar epithelial cells (Wasylnka and Moore, 2002, 2003). Although phagocytic cells are more efficient, alveolar cells can also internalize conidia, potentially leading to their destruction (Wasylnka et al., 2005; Jia et al., 2023). Some conidia may still survive, ultimately re-entering the extracellular space and contributing to persistent infection (Wasylnka and Moore, 2003; Croft et al., 2016; Culibrk et al., 2019).

In this study, we developed an *in vitro* model to investigate the interactions between *A. fumigatus* conidia and alveolar epithelial cells using micropatterning techniques. This approach allowed us to spatially confine cells on defined ECM substrates. We demonstrate that the internalization and intracellular processing of fungal conidia are significantly influenced by cell density, geometry and ECM composition. Expanding this approach to other pathogens could uncover novel mechanisms of infection and identify new targets for therapeutic intervention.

## 2 Material and Methods

### 2.1 Fungal strains and growth conditions

*A. fumigatus* DAL (wild-type, CBS 144.89) strain was cultured in filter-cap cell culture bottles on sabouraud 4 % glucose agar (15 g/L agar, 40 g/L D (+)-glucose, 10 g/L peptone, Sigma-Aldrich, 1.06404, Austria) at 37 °C for 10 days until full maturation of conidia was observed. Conidia were collected by plate flooding using sterile spore buffer (0.01 % Tween, Fisher scientific, 10113103, Germany, 0.9 % NaCl, Carl Roth, 8986.1, Germany), centrifugated at 4000 rpm for 5 minutes and resuspended in sterile spore buffer.

### 2.2 Cell culture

Human A549 cell line stably expressing Lamp1-NeonGreen (Schiefermeier-Mach et al., 2021) were cultured in a growth medium consisting of RPMI-1640 (Capricorn, RPMI-XA, Germany), 10 % fetal bovine serum (FBS, Capricorn Scientific, FBS-11A, Germany), 2 µl/ml puromycin, (Carl Roth, 0240.2, Germany) 1% penicillin/streptomycin (Capricorn Scientific, PS-B, Germany) and 1% L-glutamine (Capricorn Scientific, GLN-B, Germany).

### 2.3 Culturing cells on coverslips with micropatterns

Micropatterns were generated with deep-ultraviolet (UV) lithography on polyethylene glycol (PEG)-coated glass coverslips. Glass coverslips were incubated in a 1 mM solution of a linear PEG-silane (Rapp Polymere – reference 122000-71, Germay) in dry toluene for 20 h at 80 °C under inert atmosphere, in order to covalently immobilize a monolayer of PEG (Azioune et al., 2010; Clausen et al., 2023). The substrates were removed, rinsed intensively with isopropanol, methanol and water, and dried with nitrogen. A PEG-coated glass coverslip and a chromium-coated quartz photomask (Compugraphics, Jena) were assembled with vacuum on a holder, which was immediately exposed to deep ultraviolet (UV) light using a low-pressure mercury lamp (NIQ 60/35 XL longlife lamp, quartz tube,60 W from Heraeus Noblelight, Germany) at 5 cm distance for 5 min. Deep UV light with a wavelength of 185 nm oxidizes PEG, significantly impairs PEG antifouling properties and allows ECM protein adsorption. The patterned substrates were subsequently incubated with 150 μl of fibronectin (10 μg/ml, Sigma Aldrich, 341631, Austria) in PBS or vitronectin (3.3 µl/ml, ProSci, 91-362, USA) at 4 °C overnight and washed once with PBS. 10^5^ cells/ml (for smaller micropatterns) or 10^6^ cells/ml (for bigger micropatterns) were added and incubated in growth medium with reduced FCS (1 %) for 2 hours at 37 °C to allow adhesion. Cells were further carefully washed with warm PBS to remove unattached cells and further incubated in growth medium with reduced FCS (1 %) overnight.

### 2.4 Infection with *A. fumigatus* conidia

For infection experiments, cell were grown in 35 mm culture dishes on fibronectin- or vitronectin-coated micropatterns in serum-reduced medium and incubated for 24 hours. After the incubation time the serum-reduced medium was carefully replaced with fresh RPMI-1640 containing 10 % FBS without antibiotics. Cells were infected with 10^6^ cfu/ml *A. fumigatus* conidia. Infected cells were incubated for 1 and 3 hours at 37 °C prior fixation.

### 2.5 Fluorescence microscopy

Cells infected with *A. fumigatus* conidia were fixed as described before with sight modifications (Scheffler et al., 2014). In short, 8% paraformaldehyde (PFA, Sigma Aldrich, 104005, Austria) in PBS was slowly added directly to the cells in cell medium in proportion 1:1 and incubated for 15 minutes at 37 °C. Then, medium-PFA mix was carefully replaced with 4% PFA and incubated further 15 minutes and washed with PBS. Cells were incubated with a mixture of EasyProbe ™ ActinRed 555 to visualize actin (THP Medical Products, FP032, Austria), DAPI (Sigma-Aldrich, D9542-5MG, Austria, 1:4000) for nuclei staining and 0.05 % saponin in PBS for 1 hour at room temperature following washing in PBS and mounting on glass objective slides with Mowiol (Sigma-Aldrich, C9368, Austria).

All images were taken using IX83 Olympus Microscope (Olympus Austria) as a multiplane Z-stacks and de-convoluted using cellSens software (Olympus, deconvolution parameters: nearest neighbor 50%). 50 micropatterns from 3 biological repetitions were analyzed for each data set. Quantifications of the center of mass of fungal conidia, as well as Lamp1^+^ and a subpopulation of Lamp1^+^Actin^+^, all were saved as X, Y-coordinates, were performed in Fiji (RRID: SCR_002285). Z-stacks were examined in parallel: DAPI staining was used to count the cell number, phase contrast images were used to quantify the total number of conidia and for the identification of co-localization with Lamp1 and actin.

### 2.6 Image quantification and statistical analysis

Analysis was performed in GraphPad Prism 10.1.2 (RRID: SCR_002798, USA) and figures were prepared in Fiji and Adobe Photoshop (RRID: SCR_014199). Tests used: descriptive statistics, ordinary one-way ANOVA (multiple comparison), correlation analysis (correlation matrix, with Pearson coifficients, two-tailed, 95% confidence interval). The statistical significance of the data was determined by p-values.

## 3 Results

### 3.1 Micropattern size and cell density regulate *A. fumigatus* conidia binding and internalization

To investigate the influence of cell confinement on the internalization of *A. fumigatus* conidia, human A549 cells stably expressing Lamp1-NeonGreen were cultured on fibronectin-coated circular micropatterns of 28 µm and 60 µm in diameter. Cells were seeded at various densities to achieve 1 and 2 cells on smaller micropatterns and 10-12 cells on bigger patterns (Figure 1A). The defined cell geometry or grouping of cells on each micropattern facilitated the quantification and mapping of host-pathogen interaction events by overlaying conidia coordinates from 50 identical micropatterns. Lamp1-NeonGreen was used as a marker of conidia internalization into phagolysosomes (Schiefermeier-Mach et al., 2021). Infection of cells with dormant *A. fumigatus* conidia for 1 and 3 hours resulted in specific conidia adhesion to confined cells without binding to the PEG-coated surrounding area. The total number of detected conidia was significantly higher on big micropatterns with multicellular islands in comparison to smaller ones and strongly increased over time. However, there were no differences for 28 µm-micropattern with 1 or 2 cells (Figure 1B).

**Figure 1.**
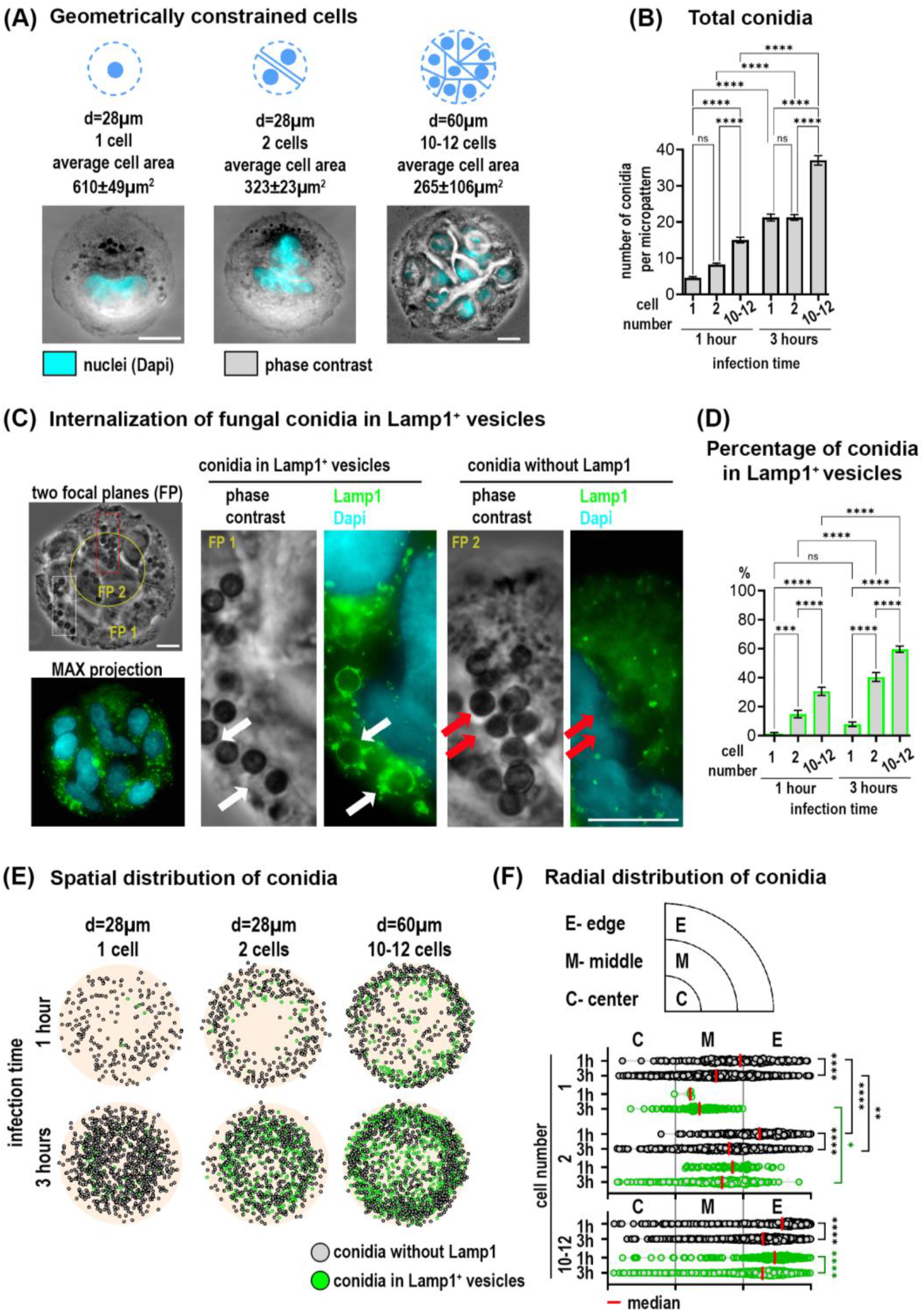
Distribution and internalization of *A. fumigatus* conidia in A549 cells constrained on circular micropatterns. (A) Representative images of cells constrained on micropatterns with diameters of 28 µm and 60 µm, and different cell densities. (B) Quantification of total conidia per micropattern at 1 and 3 hours post-infection. (C) Conidia internalization into phagolysosomes after 1 hour post-infection (Lamp1^+^ vesicles). Phase contrast image includes two focal planes superposed (FP). The overview of Lamp1 distribution is shown as the maximal Z-projection of FITC/Dapi channels of 9 focal planes (MAX projection). Magnified images show conidia in Lamp1^+^ vesicles (white arrows) and conidia without Lamp1 staining (red arrows). (D) Percentage of conidia found in Lamp1+ vesicles relative to the total conidia number at 1 and 3 hours post-infection. (E) Conidia spatial distribution map of 50 overlaid micropatterns displaying Lamp1^+^ vesicles with internalized conidia (green spheres) and conidia without Lamp1 staining (grey spheres). (F) The radial distribution of conidia defined as the distance between the micropattern center and the center of a single conidia. These distances were grouped into three segments (center, middle and edge of pattern). Depicted are conidia without Lamp1 (grey spheres) and conidia in Lamp1^+^ vesicles (green spheres). Scale bars =10 µm.

Multiplane imaging of cells by phase contrast and fluorescence microscopy revealed that some conidia were internalized into phagolysosomes (Lamp1^+^ vesicles) already after 1 hour, (Figure 1C). Since cells on micropatterns of different sizes can bind different numbers of conidia per pattern, we further normalized all values to the “total number of conidia” for each individual micropattern condition for better comparisons. Quantification of conidia in Lamp1^+^ vesicles as a part of all observed conidia showed a higher percentage in multicellular islands of 60 µm in diameter as compared to cells on 28 µm-patterns. Moreover, this percentage increased over time. Interestingly, not only the micropattern size but the number of cells per micropattern correlated with the number of phagolysosomes. There were significantly more conidia in Lamp1^+^ vesicles when 2 cells were confined to 28µm-patterns in comparison to 1 cell (Figure 1D).

Further analysis of conidia distribution revealed that binding and phagolysosomal internalization of conidia were heterogeneously distributed across the micropattern (Figure 1E). Conidia mapping and their radial distribution showed that their binding and internalization at the outer edge of the 60 µm-patterns is favored. A similar trend for 28 µm-patterns with 2 cells was also observed. The lowest number of conidia was detected in the center of all micropatterns independent of size and cell number (Figure 1E, F).

### 3.2 Cell density influences conidia trafficking in a subpopulation of Lamp1^+^Actin^+^ vesicles

To further investigate the intracellular trafficking and processing of *A. fumigatus* conidia, we additionally stained infected cells with phalloidin to detect intracellular actin distribution. We have observed that a small subset of Lamp1^+^ vesicles containing fungal conidia also displayed actin, forming a Lamp1^+^Actin^+^ subpopulation (Figure 2A). The proportion of conidia within these double-positive vesicles was relatively small compared to the overall Lamp1^+^ vesicles. However, the percentage of Lamp1^+^Actin^+^ vesicles relative to the total conidia number, showed a significant increase of these vesicles in 28 µm-patterns harboring 2 cells as compared to 1 cell or 10-12 cells on larger micropatterns (Figure 2B). Interestingly, in contrast to the population of Lamp1^+^ vesicles, which increased over time, the percentage of conidia in double-positive vesicles remained stable between 1 and 3 hours (Figure 2B). Conidia mapping further shows the number and localization of double vesicles with a tendency toward micropattern edges on bigger micropatterns.

**Figure 2.**
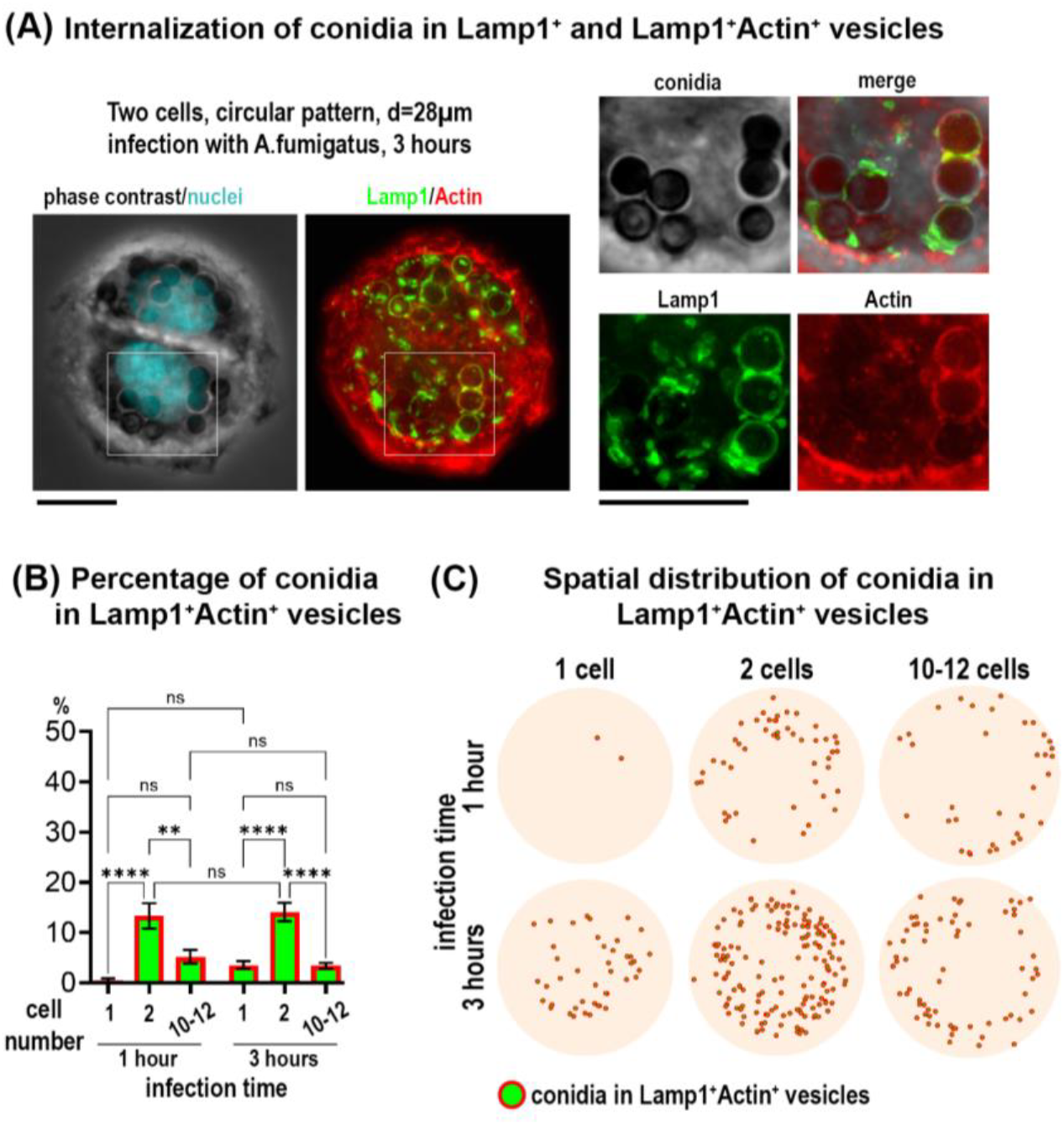
Internalization of *A. fumigatus* conidia in a subpopulation of Lamp^+^Actin^+^ vesicles in cells on circular micropatterns. (A) Representative example of two cells on circular micropatterns (28 µm). Overview of Lamp1 and Actin is shown as maximal Z-projection of FITC/TRITC channels of 9 focal planes. Magnified images show conidia in Lamp1^+^ (green) and Lamp1^+^Actin^+^ vesicles (red and green) 3 hours post-infection. Scale bars =10 µm. (B) Quantification of conidia internalized in Lamp1^+^Actin^+^ vesicles in cells constrained on circular micropatterns with different sizes and cell density at 1 and 3 hours post-infection. (C) Conidia spatial distribution map of 50 overlaid micropatterns displaying Lamp1^+^Actin^+^ vesicles with internalized conidia (green spheres in red circles).

### 3.3 ECM composition impacts conidia binding, internalization and trafficking in micropatterned cells

Next, we assessed the impact of ECM substrates in our cell model. We analyzed multicellular island constrained on 60 µm micropatterns coated with vitronectin (VN) instead of fibronectin (FN). VN is known from the literature to preferentially recruit ß3 integrins whereas FN recruits more ß1 integrins to coordinate cell adhesion (Schaufler et al., 2016). Analysis of conidia distribution on cells constricted on 60µm-patterns coated with VN, revealed that binding and phagolysosomal internalization of conidia were similarly as the ones on FN (Figure 3A,B, Figure 1E,F). With both coating conditions, more binding and internalization of conidia was observed at the outer edge of the micropattern. When we quantified the total number of conidia, we observed a decrease when using VN versus FN. However, the percentage of conidia internalized in the Lamp1^+^ vesicles did not differ from the one observed in cells micropatterned on FN (Figure 3A,B). Interestingly, there was a strong difference between the substrates in terms of the percentage of conidia internalized into the subpopulation of Lamp1^+^Actin^+^ vesicles: this percentage was increased when cells were constrained on VN.

**Figure 3.**
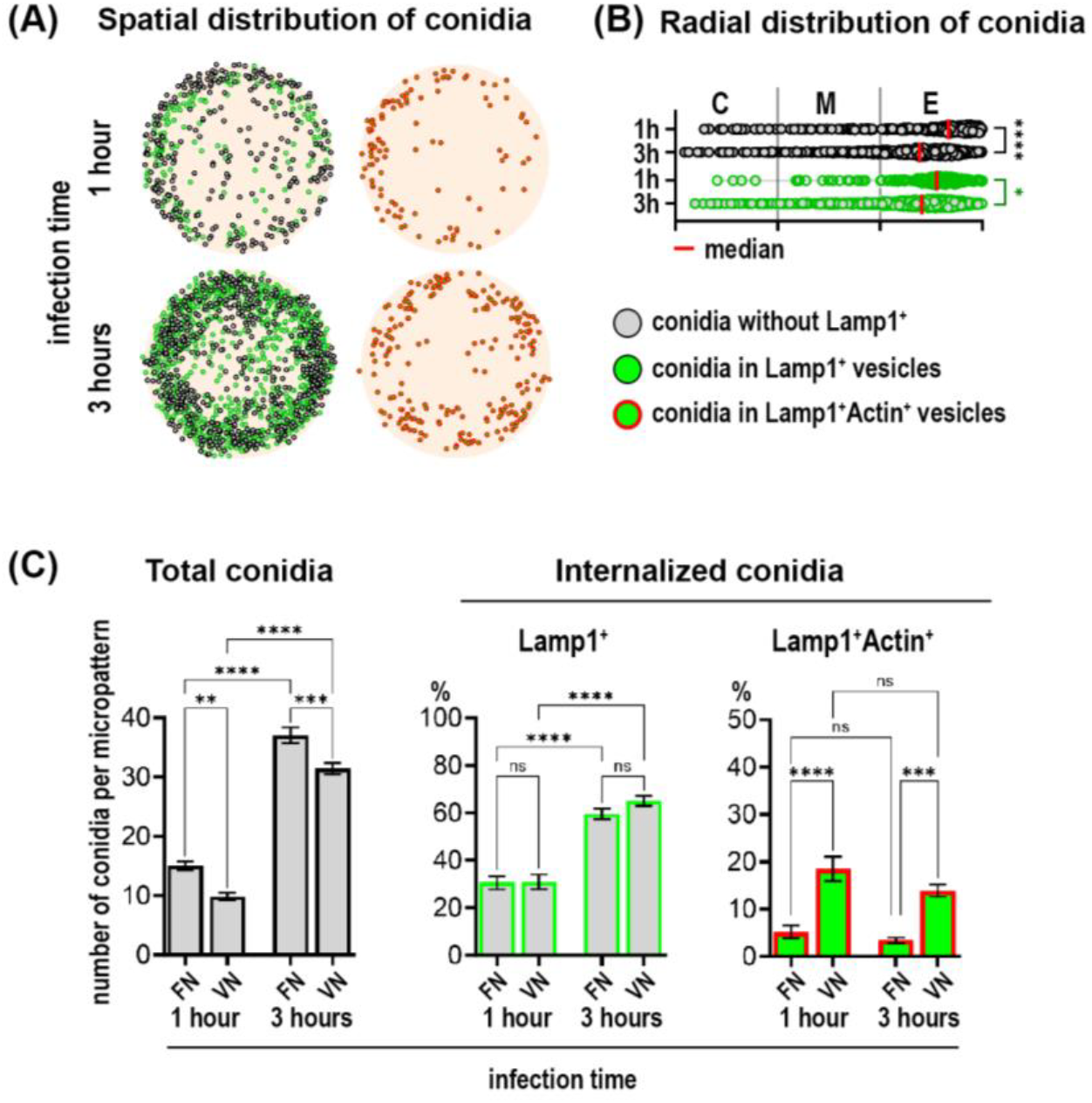
ECM composition affects conidia internalization. (A) Conidia spatial distribution map of 50 overlaid micropatterns displaying conidia without Lamp1 (grey spheres), conidia in Lamp1^+^ vesicles (green spheres) and conidia in the subpopulation of Lamp1^+^Actin^+^ vesicles (green spheres in red circles). (B) The radial distribution of conidia defined as the distance between the micropattern center and the center of a single conidia. These distances were grouped into three segments (center, middle and edge of pattern). Depicted are conidia without Lamp1 (grey spheres) and conidia in Lamp1^+^ vesicles (green spheres). (C) Quantification of total conidia number, conidia internalized in Lamp1^+^ and in Lamp1^+^Actin^+^ vesicles at 1 and 3 hours post-infection.

### 3.4 Cell shape and cell number impact conidia distribution and internalization in cells constricted on X-shaped micropatterns

To further investigate the impact of cell geometry on the cell-conidia interactions, we seeded A549-Lamp1-NeonGreen cells on X-shaped patterns with a diagonal of 28 µm. By varying the number of cells, we established 3 distinct cell densities: 1, 2 and 4 cells per micropattern, each resulting in characteristic changes in cell morphology due to the shape constraints imposed by the X-pattern (Figure 4A).

**Figure 4.**
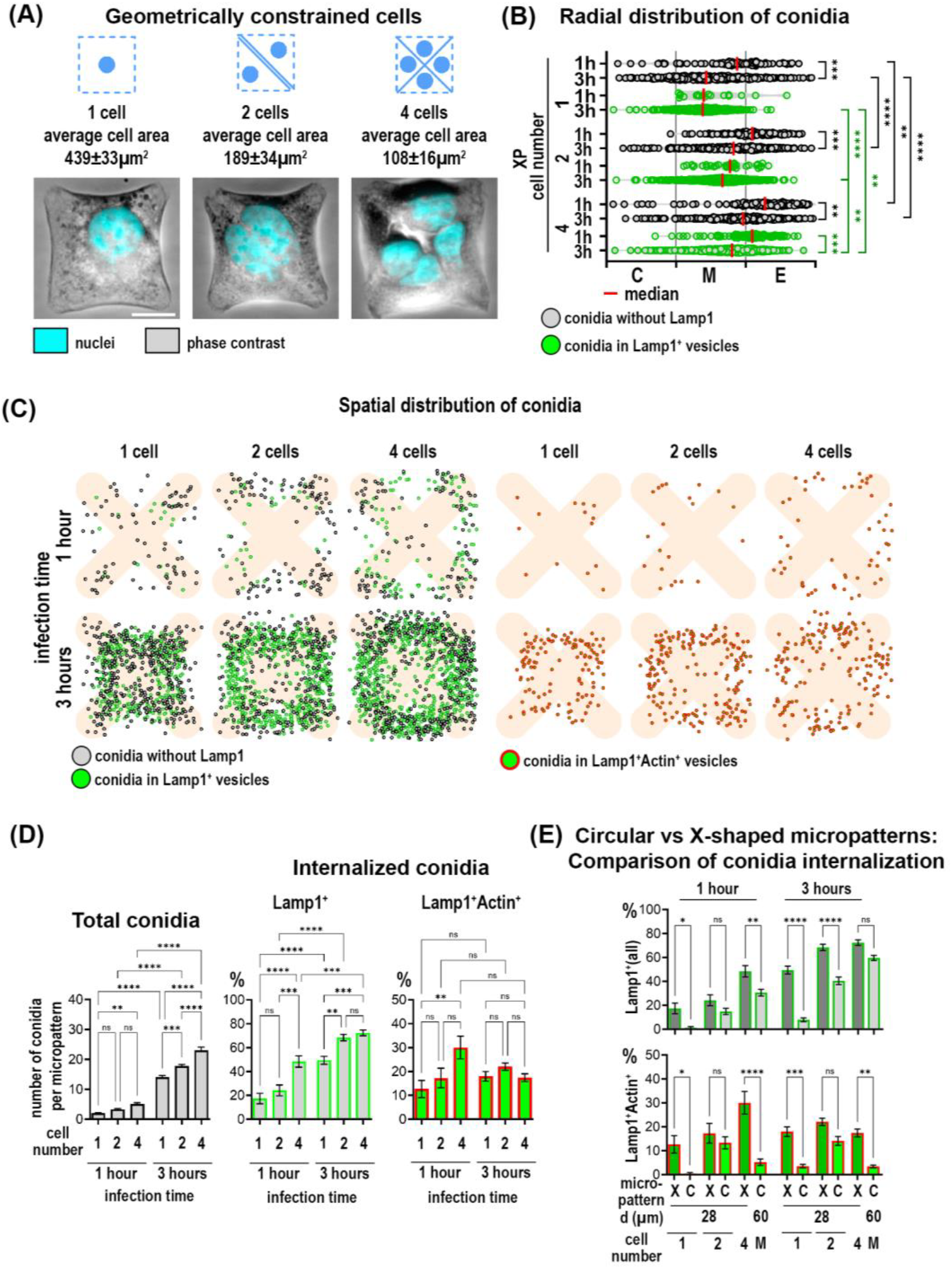
Distribution and internalization of *A. fumigatus* conidia in A549 cells constrained on X-shaped micropatterns. (A) Representative phase contrast images of cells constrained on X-micropatterns with different cell densities. Scale bars =10 µm. (B) The radial distribution of conidia defined as the distance between the micropattern center and the center of a single conidia. These distances were grouped into three segments (center, middle and edge of pattern). Depicted are conidia without Lamp1 (grey spheres) and conidia in Lamp1^+^ vesicles (green spheres). (C) Map of conidia spatial distribution in 50 overlaid micropatterns showing Lamp1^+^ vesicles with internalized conidia (green spheres), conidia without Lamp1 staining (grey spheres) and conidia in Lamp1^+^Actin^+^ vesicles. (D) Quantification of total conidia number, conidia internalized in Lamp1^+^ and in Lamp1^+^Actin^+^ vesicles at 1 and 3 hours post-infection.

Upon infection with *A. fumigatus* conidia for 1 and 3 hours, we observed notable differences in conidia internalization across these conditions. After 1 hour of infection, while the total number of conidia did not differ, there was an increased percentage of conidia in Lamp1^+^ vesicles in the 4-cell configuration in comparison to 1 cell and 2 cells. After 3 hours of infection, the total number of conidia correlated with cell number was the highest in the 4-cell micropattern. Internalization of conidia in Lamp1^+^ vesicles increased after 3 hours compared to 1 hour and was higher in micropatterns with 2 and 4 cells compared to those with only 1 cell. The percentage of conidia in double Lamp1^+^Actin^+^ vesicles did not differ significantly between the X-micropatterns with different cell numbers and did not change over time.

Interestingly, there were notable differences when comparing X-shaped to circular micropatterns (Figure 4E). Conidia were internalized more efficiently by cells constrained on X-micropatterns. Moreover, the percentage of double Lamp1^+^Actin^+^ vesicles was also higher in these cells (Figure 4E). To further elucidate the relationship between different parameters and conidia internalization, we performed a correlation analysis using a correlation matrix and Pearson correlation coefficients. This analysis revealed that the percentage of conidia in Lamp1^+^ and Lamp1^+^Actin^+^ vesicles was negatively correlated with cell area at both 1 hour and 3 hours. This negative correlation was moderate after 1 hour (r=-0.447) and strong after 3 hours (r=-0.716). The correlation of cell area with Lamp1^+^Actin^+^ vesicles was negative and moderate both after 1 hour (r=-0.310) and 3 hours (r=-0.299). No other significant correlations with other parameters were found.

## 4 Discussion

Our study emphasizes the role of biological boundary conditions in the regulation of host-pathogen interactions in the lung. The complex geometry of the alveoli and the mechanical forces during respiration establish a unique environment that influences cellular behavior (Roan and Waters, 2011). Previous studies showed that cell geometry, ECM composition, and mechanical forces affect essential lung functions, including alveolar permeability (Cavanaugh et al., 2005), surfactant secretion (Edwards, 2001), and apoptosis (Hammerschmidt et al., 2007). Here, we extend this understanding by demonstrating that boundary conditions also regulate the binding, internalization and intracellular fate of *A. fumigatus* conidia in alveolar epithelial cells. Our data provided new insights into how the lung’s microenvironment can influence infection outcomes, particularly under conditions of altered mechanical and biochemical cues.

The choice of micropatterns was particularly advantageous for our study as it allowed precise control over cell geometry, density, and ECM composition, simulating specific aspects of the alveolar environment. We observed that constraining cells on micropatterns strongly increased the efficiency of conidia binding and phagolysosomal internalization compared to previous studies with no cell confinement (Wasylnka and Moore, 2002; Jia et al., 2023), reaching up to 30 % already after one hour of infection. Our results further showed that cell shape and density had profound effects on conidia uptake. Cells on X-shaped micropatterns demonstrated higher internalization efficiency compared to circular ones, suggesting that specific geometric configurations create environments more conducive to pathogen internalization. Previous studies in single cells showed that the size and shape of the micropattern influence contractility and actin organization (Pitaval et al., 2010; Théry, 2010). Cells on X-shaped micropatterns had regions of localized higher tension, since this shape forced them to stretch along the diagonals, increasing mechanical stress at intersections and edges (James et al., 2008). In circular patterns, cellular tension was more evenly distributed, but still higher at the periphery than the center (Théry et al., 2006; Théry and Bornens, 2006). Such control over cell mechanics and geometry is a significant benefit of using micropatterning techniques, enabling detailed investigation into how specific physical constraints can enhance or limit pathogen internalization.

In multicellular islands, micropatterns dictate not only individual cell behavior but also the collective dynamics of cell groups (Théry, 2010). Peripheral cells were observed to be more contractile compared to those in the center, with higher traction forces at the edges. This variation arose due to spatial confinement, leading to differences in cell proliferation and differentiation across the micropattern (Nelson et al., 2005; He et al., 2015; Lin et al., 2022; Nelson et al., 2024). Although larger single cells might theoretically offer more surface area for conidia binding, our results showed that smaller cells within multicellular islands internalized more conidia than larger, single cells. This was significant on both circular and X-shaped micropatterns. The geometric constraint affected the recruitment of actin to the phagolysosomal membranes. Cells on X-shaped micropatterns exhibited a higher frequency of Lamp1^+^Actin^+^ vesicles compared to circular ones. Interestingly, while the percentage of conidia in Lamp1^+^ vesicles increased over the infection period, the percentage of Lamp1^+^Actin^+^ vesicles plateaued after one hour, suggesting that actin recruitment to phagolysosomes may be limited to specific stages of vesicle maturation.

Moreover, the binding and internalization events were more frequent at the periphery of the multicellular micropattern. This finding was consistent with the work of Feng et al (Feng et al., 2023), who showed that spatial tension heterogeneity leads to uneven collagen expression and increases bacterial adhesion at the periphery of micropatterned multicellular islands. Our results extended these observations to interactions with fungal pathogens. This phenomenon may be particularly relevant in the lung, where mechanical heterogeneity arises from alveolar geometry and the forces generated during respiration (Roan and Waters, 2011). In pathological conditions such as fibrosis or emphysema, where ECM stiffness is altered, these effects may be enhanced, potentially creating regions more susceptible to infection (Colebatch et al., 1973; Wilson, 1983).

Our findings also highlighted the influence of ECM composition on conidia internalization and trafficking. By varying coating substrates, distinct effects on conidia processing were observed. When cells were plated on vitronectin (VN), the integrin activation profile shifted from α5β1 (a main integrin for fibronectin (FN)) to αvβ3 integrin (Schaufler et al., 2016). This shift decreased overall conidia binding compared to FN-coating but increased the proportion of conidia internalized into Lamp1^+^Actin^+^ vesicles. These results suggested that integrin-mediated signaling pathways play a role in determining the fate of internalized conidia. It aligns with previous studies showing that different integrin receptors, specifically α5β1 and αvβ3, influence cytoskeletal dynamics, vesicular trafficking and phagocytosis (Dupuy and Caron, 2008; Bastounis et al., 2022). The increased presence of Lamp1^+^Actin^+^ vesicles in cells on VN substrate indicated that β3 integrin engagement may favor a trafficking pathway involving dynamic actin rearrangement, potentially facilitating more efficient phagosome maturation.

In summary, our study emphasized the significance of biological boundary conditions in shaping host-pathogen interactions. The heterogeneous internalization observed across different micropattern configurations suggested that some lung regions may be more prone to infection than others, especially during chronic disease states. This highlighted the importance of spatially localized mechanical cues in determining the efficiency of conidia uptake and processing, with potential implications for understanding fungal infection establishment in the lung. Future research should aim to dissect the molecular mechanisms linking mechanical forces to pathogen internalization, particularly under varying disease conditions that alter ECM composition and stiffness or respiratory mechanics.

## 5 Conflict of Interest

*The authors declare that the research was conducted in the absence of any commercial or financial relationships that could be construed as a potential conflict of interest*.

## 6 Author Contributions

The Author Contributions are: writing: NSM, JP, LH, LL, OV, SP; conceptualization: NSM, JP; formal analysis: LH, LL, OV; methodology: JP, LH, LL; project administration: NSM, PJ, LH; resources: NSM, JP, SP, supervision: NSM; validation: JP, LH; visualization: NSM, OV, LH, LL; funding acquisition: NSM, LL.

## 7 Funding

This research was funded by Tirol Wissenschaftsfoerderung TWF to Natalia Schiefermeier-Mach (F.33484) and to Lukas Lechner (F.47867).

## 8 Data Availability Statement

The datasets are available upon request.

## Notes

### Competing Interest Statement

The authors have declared no competing interest.

